# Residual Structure of Unfolded Ubiquitin as Revealed by Hydrogen/Deuterium-Exchange 2D NMR

**DOI:** 10.1101/429167

**Authors:** M. S. Chandak, T. Nakamura, M. Yagi-Utsumi, T. Yamaguchi, K. Kato, K. Kuwajima

## Abstract

The characterization of residual structures persistent in unfolded proteins in concentrated denaturant solution is currently an important issue in studies of protein folding, because the residual structure present, if any, in the unfolded state may form a folding initiation site and guide the subsequent folding reactions. Here, we thus studied the hydrogen/deuterium (H/D)-exchange behavior of unfolded ubiquitin in 6.0 M guanidinium chloride at pH 2.6 and 20°C. We employed a dimethylsulfoxide (DMSO)-quenched H/D-exchange NMR technique with the use of spin desalting columns, which allowed us to make a quick medium exchange from 6.0 M guanidinium chloride to a quenching DMSO solution. The technique is particularly effective for studies of the H/D-exchange kinetics of unfolded proteins in concentrated denaturant. By the backbone resonance assignment of the hetero-nuclear single quantum coherence spectrum of ^15^N-labeled ubiquitin in the DMSO solution, we successfully investigated the H/D-exchange kinetics of 27 identified peptide amide groups in the ubiquitin sequence. Although most of these amide groups were not protected, the four amide groups of Ile3, Val5, Ile13 and Leu73 were weakly but significantly protected with a protection factor of 2.5–3.0, indicating that there were residual structures in unfolded ubiquitin and that these amide groups were 60–67% hydrogen-bonded by the residual structures. We show that the first native β-hairpin, composed of residues 2–16 in the native ubiquitin structure, is partially structured even in 6.0 M guanidinium chloride and that the amide group of Leu73 is protected by a nonnative hydrogen-bonding interaction. From comparison with the previous folding studies of ubiquitin, it is concluded that the residual native β-hairpin in unfolded ubiquitin forms a folding initiation site and guides the subsequent folding reactions of the protein.

## Introduction

For many years, globular proteins in concentrated denaturant solution (6 M guanidinium chloride (GdmCl) or 8 M urea) were assumed to be in the fully unfolded random-coil state (1, 2). However, more recently, evidence for the persistence of residual structures in unfolded proteins in a concentrated denaturant has been reported for several different proteins (3–7). Therefore, in order to elucidate the molecular mechanisms of protein folding, characterization of the residual structures present in the unfolded state is important (8). The initial starting state in kinetic refolding experiments is often the unfolded state in a concentrated denaturant, and hence, the persistent residual structure present, if any, in the unfolded state may form a folding initiation site and guide the subsequent folding reactions.

The hydrogen/deuterium (H/D)-exchange method combined with 2D NMR spectroscopy is a powerful technique to detect and characterize the residual structures of an unfolded protein, and the NMR signals of peptide amide (NH) groups may be monitored by ^1^H–^15^N hetero-nuclear single quantum coherence (HSQC) spectra of the ^15^N-labeled protein. However, the observed exchange rates of the NH groups are often too fast to follow directly by 2D NMR. Therefore, rapid quenching of the H/D-exchange reaction is required for studying the H/D-exchange kinetics of the unfolded protein in a concentrated denaturant. Rapid refolding to the native state is an effective approach for the rapid quenching, and the rapid refolding technique has been used for studying the unfolded-state H/D-exchange kinetics of hen egg-white lysozyme in 8 M urea at pH 2.0 (9) and human fibroblast growth factor 1 in 4 M urea at pH 4.0 (10). However, a drawback of the use of rapid refolding for quenching the H/D exchange is that we can detect and monitor only the stably protected NH protons in the native state, and hence, it is impossible to detect NH protons that are unprotected in the native state.

In our previous study, we reported that the use of spin desalting columns in dimethylsulfoxide (DMSO)-quenched H/D-exchange NMR (^1^H–^15^N HSQC) experiments is very useful for investigating fast-exchanging NH protons in the unfolded protein in a concentrated denaturant (11). We can realize a quick exchange of the medium from the concentrated denaturant to a quenching DMSO solution by the use of a spin desalting column, and analyze the signals of the remaining NH protons of the protein in the DMSO solution. We applied this method for studying the H/D-exchange kinetics of unfolded ubiquitin in 6.0 M GdmCl at pH* 2.6 and 20°C; pH* indicates the pH meter reading. Although we successfully observed the exchange kinetics of a number of the NH protons, the assignment of the NH proton signals in the ^1^H–^15^N HSQC spectrum was not available in the previous study (11).

To investigate the H/D-exchange kinetics of individually identified NH proton signals of ubiquitin, here, we carried out the backbone assignment of the HSQC spectrum of the protein in the DMSO solution by the use of a ^1^H 920-MHz NMR instrument. As a result, we could successfully investigate the H/D-exchange kinetics of 27 NH groups in the ubiquitin sequence. Although most of these NH groups were fully exposed to solvent without any significant protection against H/D exchange, the four NH groups of Ile3, Val5, Ile13 and Leu73 were weakly but significantly protected with a protection factor of 2.5–3.0, indicating that there were residual structures in unfolded ubiquitin and that these NH groups were 60–67% hydrogen (H)-bonded in 6.0 M GdmCl. We show that the first native β-hairpin, composed of residues 2–16 of ubiquitin in the native structure (12), is partially structured even in 6.0 M GdmCl and that the NH group of Leu73 is protected by a nonnative H-boding interaction in the unfolded state. From comparison of the present results with the previous studies on the folding kinetics of ubiquitin (13–16), it is concluded that the residual native β-hairpin in unfolded ubiquitin forms a folding initiation site and guides the subsequent folding reactions of the protein.

## Material and Methods

### Materials

DMSO-*d*_6_ and D_2_O were purchased from Cambridge Isotope Laboratories Inc. (Andover, MA). Dichloroacetic acid was purchased from Tokyo Chemical Industry Co., Ltd. (Tokyo, Japan), and used for pH adjustment of a quenching DMSO solution (95% DMSO-*d*_6_/5% D_2_O, pH* 5.0). GdmCl was purchased from Nakalai Tesque Inc. (Kyoto, Japan). Deuterated GdmCl was produced by repeated cycles of dissolution of GdmCl in D_2_O followed by lyophilization.^15^N-labeled and {^13^C,^15^N}-double-labeled human ubiquitin were expressed in *Escherichia coli* host cells BL21-Codon-Plus or BL21(DE3) at 37°C in M9 minimum medium as recombinant proteins and purified as described previously (11, 17). Ubiquitin thus obtained was flanked by an N-terminal hexahistidine-tag sequence derived from a cloning vector pET28a (Novagen) used for the protein expression (see Fig. 1).

**FIGURE 1.**
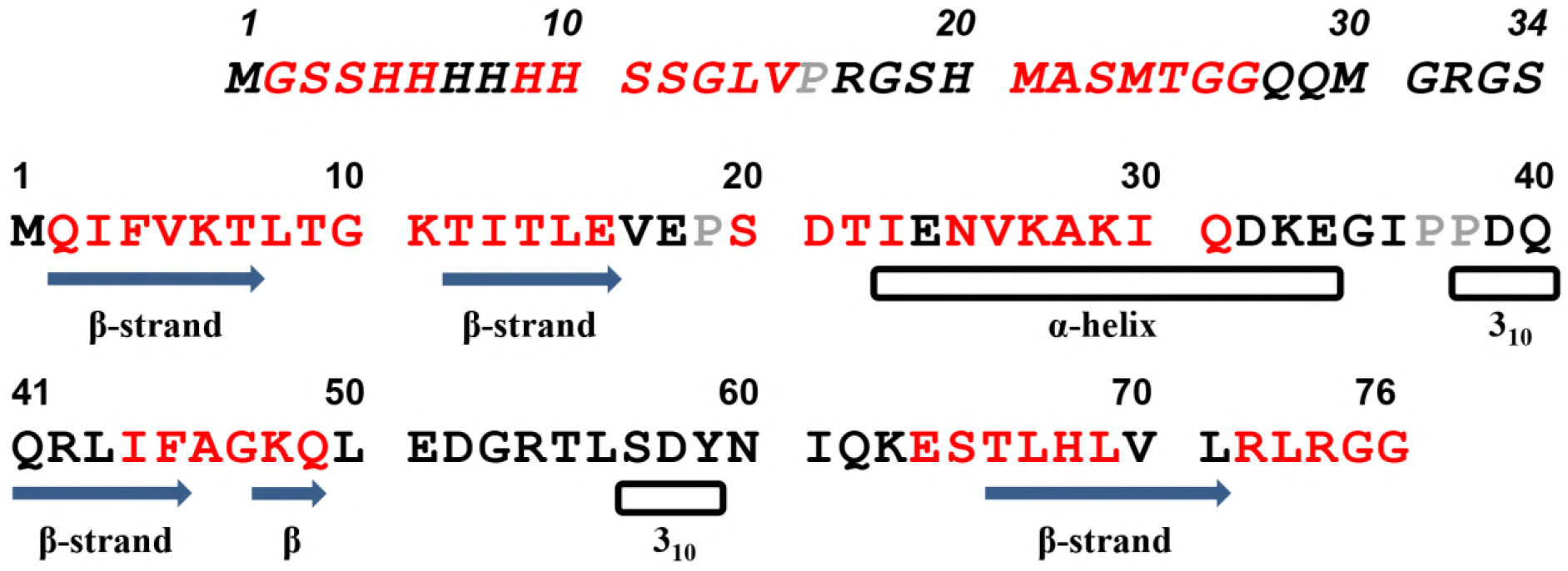
The primary structure of the protein used in this study. The first 34 residues in italics are the N-terminal extra residues, and the next 76 residues in roman type are the residues of human ubiquitin. The residues in the red one-letter code are those whose peptide NH signals in the ^1^H–^15^N HSQC spectrum of the 15N-labeled protein have been assigned. The locations of the secondary structures in native ubiquitin (PDB code: 1UBQ) are shown by arrows (β-strands) and open rectangles (helices).

### DMSO-quenched H/D-exchange experiments

The H/D-exchange reaction of unfolded ubiquitin was started by 10-fold dilutions of 3 mM ^15^N-labeled ubiquitin unfolded in 6.0 M GdmCl in H2O into 6.0 M deuterated GdmCl in D_2_O at pH* 2.6 and 20.0°C. At each pre-determined exchange time, 1.0 mL of the reaction mixture was taken, the reaction was quenched in liquid nitrogen, and the frozen mixture was kept at −85°C until the medium exchange and the subsequent NMR measurement. For the NMR measurement, the frozen sample was thawed at room temperature, the medium containing 6.0 M GdmCl was exchanged for the quenching DMSO solution by using a spin desalting column (Zeba^TM^ Spin Desalting Column 89891; Thermo Scientific), and the ^1^H–^15^N HSQC spectrum of the protein was measured by a ^1^H 500-MHz NMR instrument. The use of spin desalting columns for the medium exchange in the DMSO-quenched H/D-exchange experiments is superior to conventionally used refolding or lyophilization. The details of the use of spin desalting columns and the typical HSQC spectra thus obtained with different H/D-exchange times were reported previously (11).

### NMR measurements

The standard ^1^H–^15^N HSQC spectra of the protein in the DMSO solution with different H/D-exchange times were acquired at 25°C on a Bruker Avance 500 spectrometer. To achieve the backbone resonance assignment of the protein in the DMSO solution, three-dimensional CBCACONH and HNCACB experiments were performed on a JEOL ECA 920-MHz spectrophotometer at 25°C. All NMR data were processed using NMRPipe (18) and NMRView (19).

### Data analysis

The observed kinetic exchange curves, given by the volumes [*Y*(*t*)] of cross-peaks in the ^1^H–^15^N HSQC spectra as a function of the H/D-exchange time (*t*), were single exponential under the H/D-exchange condition (6.0 M GdmCl, 90% D_2_O/10% H2O at pH* 2.6 and 20.0°C), and fitted to the equation 
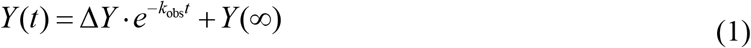
where Δ*Y, k*_obs_ and *Y*(∞) are the kinetic amplitude, the observed H/D-exchange rate constant, and the final value of the peak volume, respectively. The fitting was performed using the IGOR Pro 6.37 software package (WaveMetrix). The predicted intrinsic rate constant (*k*_int_) of the H/D exchange for the non-protected NH protons was calculated by the methods of Bai *et al*. (20) and Connelly *et al*. (21). We used the program SPHERE for the calculation of *k*_int_; SPHERE is accessible through the internet at the following URL, http://www.fccc.edu/research/labs/roder/sphere/sphere.html.

## Results

### The primary structure

Figure 1 shows the primary structure of ubiquitin used in this study. The protein had an N terminal extension of 34 residues, which derived from a linker sequence of a His-tag cloning vector used for the protein expression. These extra residues are shown in italics in Fig. 1. The 76 residues from residue 35 (Met) to residue 110 (Gly) correspond to the residues of human ubiquitin, and these residues are re-numbered from 1 to 76 following the residue number of ubiquitin. To distinguish these residues from the 34 extra N-terminal residues, the ubiquitin residues are shown in the roman type. The three-dimensional structure of native ubiquitin is composed of three helices (a 12-residue α-helix and two one-turn 310-helices) and five β-strands (12), and the locations of these secondary structures, determined by the DSSP program (22), are also shown in Fig. 1.

### Backbone assignment of the HSQC spectrum

To make the backbone assignment of the HSQC spectrum of ubiquitin in the DMSO solution, we carried out three-dimensional CBCACONH and CBCANH experiments of {^13^C, ^15^N}-double-labeled ubiquitin recorded on the ^1^H 920-MHz NMR instrument. The protein has 105 peptide NH groups (32 in the N-terminal extension and 73 in the ubiquitin sequence), and we identified 62 out of the 105 NH groups or 43 out of the 73 NH groups in the ubiquitin sequence. Figure 2 shows a HSQC spectrum of the uniformly ^15^N-labeled protein thus obtained with backbone assignment, where the ubiquitin residues are shown in roman type (red) and the N-terminal extra residues are shown in underlined italics (cyan). In Fig. 1, the 62 residues with backbone assignment are shown with a red one-letter code.

**FIGURE 2.**
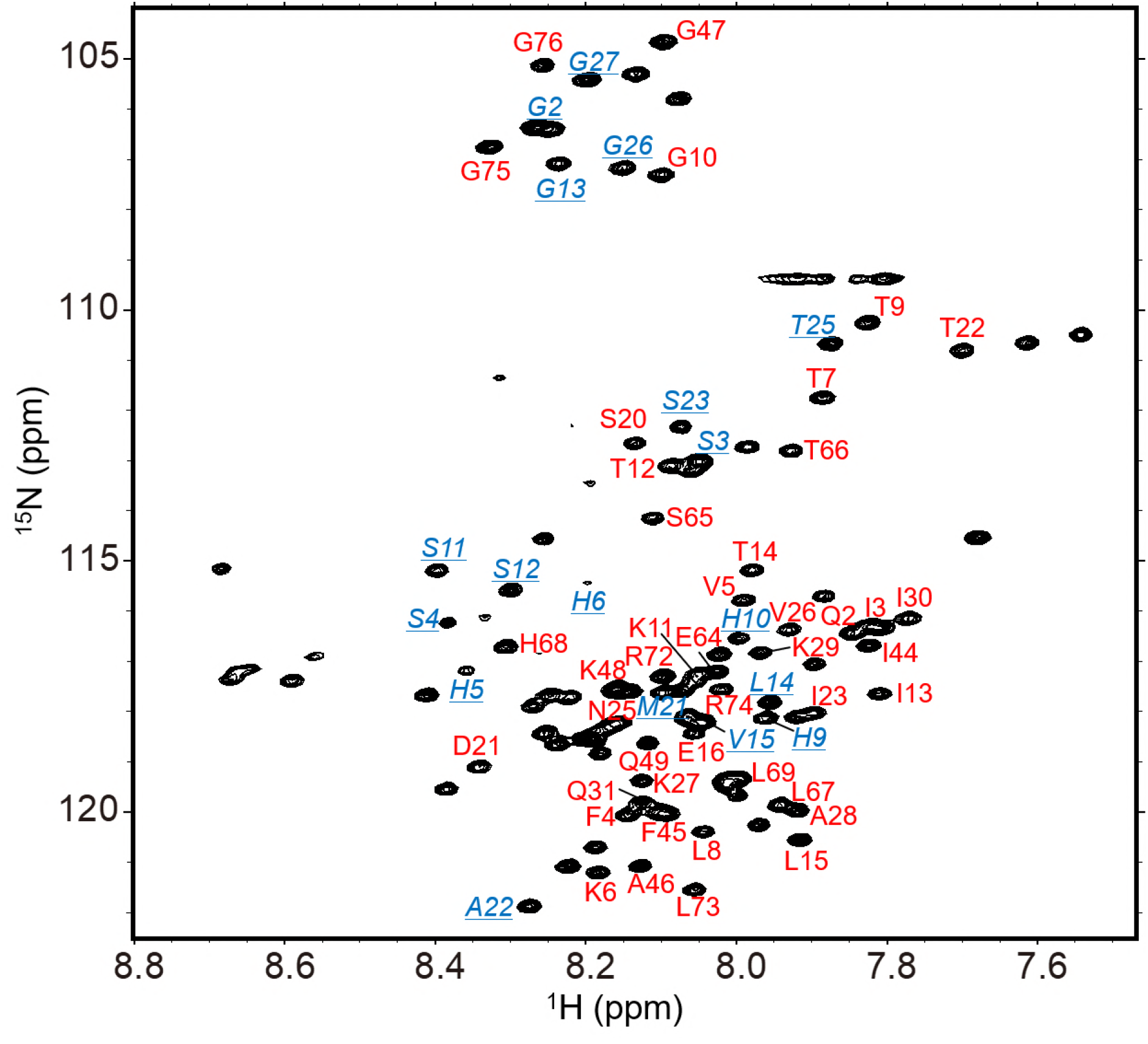
The ^1^H–^15^N HSQC spectrum of uniformly ^15^N-labeled ubiquitin in the DMSO solution (95% DMSO-*d*_6_/5% D_2_O, pH* 5.0) recorded on the ^1^H 920-MHz NMR instrument at 25°C. The resonance assignment for the ubiquitin sequence is shown in roman type (red), and the assignment for the N-terminal extra residues is shown in underlined italics (cyan).

### The H/D-exchange kinetics of individual NH groups

The protein solutions with different H/D-exchange times were exchange-quenched by the DMSO solution, and their HSQC spectra were recorded by ^1^H 500-MHz NMR; for typical HSQC spectra thus obtained, see (11). On the basis of the backbone assignment of the ^1^H– ^15^N HSQC spectrum (Fig. 2), we investigated the H/D-exchange kinetics of the individual NH groups whose cross peaks in the HSQC spectra were assigned. The exchange kinetics were given by the peak volumes of cross peaks in the HSQC spectra as a function of the exchange time. By excluding NH groups whose cross peaks were extensively overlapping with each other or too weak in intensity to follow the exact kinetics, we successfully followed the H/D-exchange kinetics of 32 NH groups. All the exchange kinetics were well represented by a single-exponential function. Figure 3 shows typical H/D-exchange curves for four residues, Val5, Thr9, Lys48 and Thr66, and the values of the observed exchange rate constant (*k*_obs_) for the 32 NH groups are summarized in Table 1.

**FIGURE 3.**
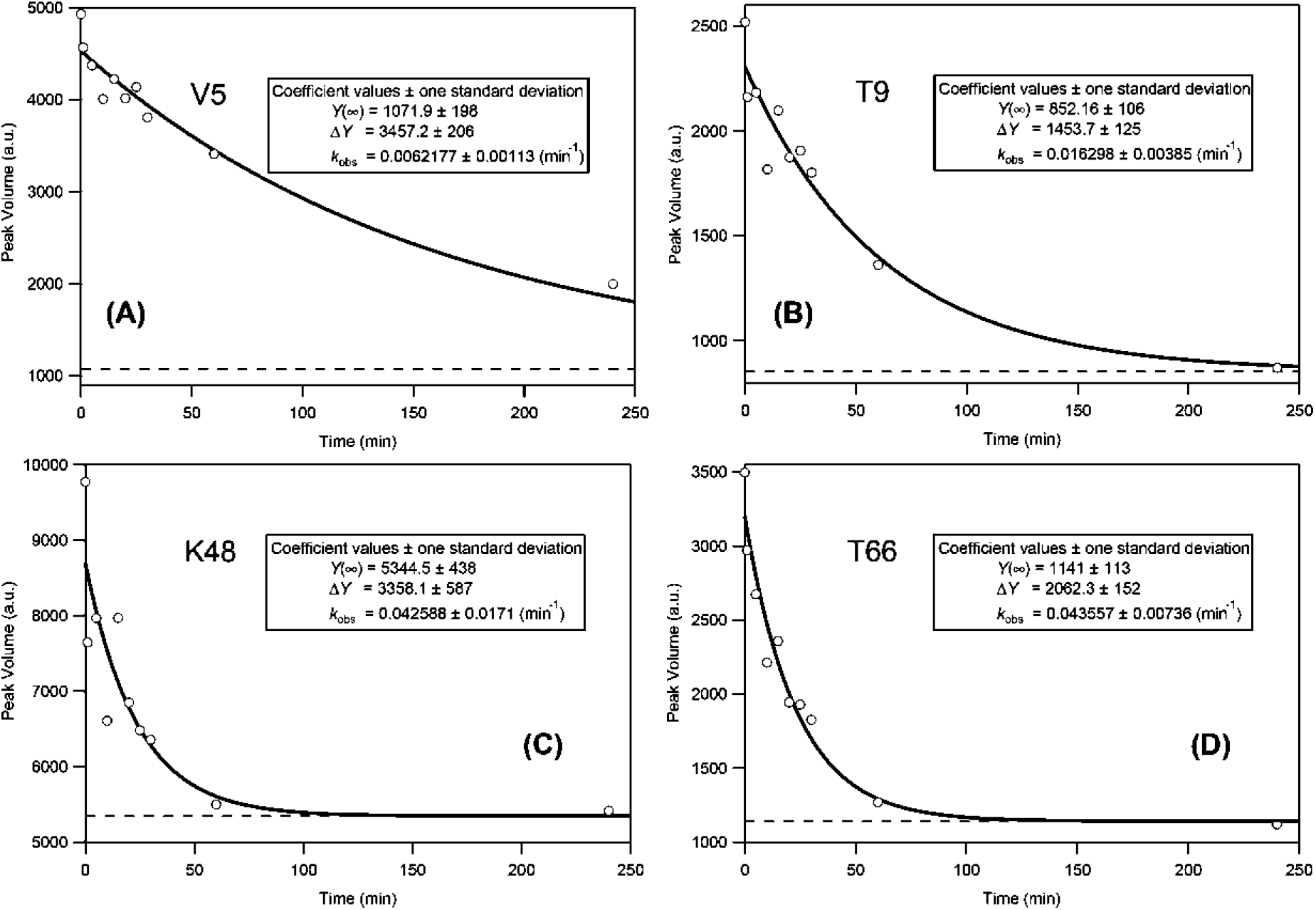
The H/D-exchange curves for Val5 (A), Thr9 (B), Lys48 (C) and Thr66 (D). The solid lines are the theoretical curves best-fitted to a single-exponential function (Eq. 1). The broken lines indicate the peak volumes after a complete exchange (i.e., *Y*(∞) in Eq. 1). The *k*_obs_ values for the four residues are as follows: (A) (6.2 ± 1.1)×10^−^3 min^−^1; (B) (1.6 ± 0.4)×10^−^2 min^−^1; (C) (4.3 ± 1.7)×10^−^2 min^−^1; and (D) (4.4 ± 0.7)×10^−^2 min^−^1.

**Table 1.** The H/D-exchange Parameters of the Peptide NH Groups of Unfolded Ubiquitin at 6.0 M GdmCl (pH* 2.6 and 20°C) and the Hydrogen (H)-bond Lengths and Acceptors in the Native Structure (PDB Code: 1UBQ)

| Residue | $k_{\text{obs}}$ ($\text{min}^{-1}$) | $k_{\text{int}}$ ($\text{min}^{-1}$) | $P$ | H-bond length ($\text{\AA}$)<sup>a</sup> | Acceptor Residue<sup>b</sup> |
| --- | --- | --- | --- | --- | --- |
| <u>S3</u> | $3.6 \pm 2.1 \times 10^{-2}$ | $4.56 \times 10^{-2}$ | 1.3 | — | — |
| <u>L14</u> | $2.7 \pm 1.1$ | 2.71 | 1.0 | — | — |
| <u>S23</u> | $4.8 \pm 2.1$ | 3.51 | 0.7 | — | — |
| <u>T25</u> | $2.8 \pm 0.3$ | 2.31 | 0.8 | — | — |
| <u>G26</u> | $4.8 \pm 2.5$ | 3.48 | 0.7 | — | — |
| I3 | $0.76 \pm 0.17$ | 1.87 | 2.5 | 3.05 | L15 |
| F4 | $2.0 \pm 0.5$ | 1.91 | 1.0 | 2.86 | S65 |
| V5 | $0.62 \pm 0.11$ | 1.84 | 3.0 | 2.79 | I13 |
| K6 | $1.8 \pm 0.6$ | 2.20 | 1.2 | 2.90 | L67 |
| T7 | $1.6 \pm 0.7$ | 2.32 | 1.4 | 2.91 | K11 |
| L8 | $1.4 \pm 0.4$ | 1.98 | 1.4 | | |
| T9 | $1.6 \pm 0.4$ | 2.09 | 1.3 | | |
| G10 | $3.9 \pm 3.2$ | 3.48 | 0.9 | | |
| K11 | $3.9 \pm 1.5$ | 3.21 | 0.8 | | |
| T12 | $2.7 \pm 0.8$ | 2.32 | 0.9 | | |
| I13 | $0.73 \pm 0.18$ | 1.82 | 2.5 | 2.72 | V5 |
| T14 | $1.5 \pm 0.3$ | 1.92 | 1.3 | | |
| L15 | $1.9 \pm 1.4$ | 1.98 | 1.0 | 3.05 | I3 |
| S20 | $6.7 \pm 4.2$ | 2.75 | 0.4 | | |
| T22 | $4.7 \pm 1.4$ | 3.45 | 0.7 | | |
| I23 | $1.5 \pm 0.4$ | 1.82 | 1.2 | 2.86 | R54 |
| N25 | $3.8 \pm 1.2$ | 5.55 | 1.5 | | |
| V26 | $1.6 \pm 0.3$ | 2.07 | 1.3 | 3.12 | T22 |
| K29 | $2.4 \pm 1.0$ | 2.61 | 1.1 | 2.93 | N25 |
| A46 | $2.6 \pm 1.5$ | 2.93 | 1.1 | | |
| G47 | $3.6 \pm 3.6$ | 3.78 | 1.0 | | |
| K48 | $4.3 \pm 1.7$ | 3.21 | 0.7 | 2.84 | F45 |
| Q49 | $2.6 \pm 1.1$ | 2.70 | 1.0 | | |
| T66 | $4.4 \pm 0.7$ | 2.57 | 0.6 | | |
| L69 | $1.4 \pm 0.4$ | 2.64 | 1.9 | 2.93 | K6 |
| L73 | $0.84 \pm 0.26$ | 2.07 | 2.5 | | |
| G75 | $7.0 \pm 6.4$ | 3.72 | 0.5 | | |

<sup>a</sup>The H-bond distance between the peptide NH group of the residue in the first column and the peptide carbonyl (CO) group of the acceptor residue in native ubiquitin is listed.
<sup>b</sup>The peptide CO group of the acceptor residue forms an H bond with the NH group of the residue in the first column. Blank cells indicate the absence of the acceptor CO group.

### Intrinsic H/D-exchange rate constants and protection factors

The intrinsic rate constant (*k*_int_) for the chemical H/D-exchange reaction of an NH group in a fully randomly coiled polypeptide is sensitive to neighboring residues, solution conditions (pH and temperature) and equilibrium and kinetic isotope effects, and methods for predicting the kint value of a given NH group have been reported by Bai *et al*. (20) and Connelly *et al*. (21). We used these methods to evaluate the *k*_int_ values of the individual peptide NH groups of the protein. However, Loftus *et al*. (23) previously reported that the presence of 6.0 M GdmCl resulted in a two-fold deceleration of the H/D exchange rate of the peptide NH groups in the acid-catalyzed pH region at 20°C. Therefore, the *k*_int_ values listed in Table 1 are those after the correction for the GdmCl effect, i.e., *k*int = (the exchange rate constant predicted by the above methods)/2. The protection factor (*P*) that is the degree of protection of a given NH group against the H/D exchange with a solvent deuteron, is given by: 
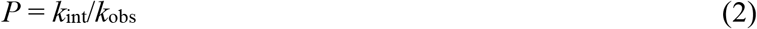

The *P* values thus obtained for the peptide NH groups of the protein are also listed in Table 1.

From Table 1, most NH groups have a *P* value close to unity (i.e., 1.0 ± 0.4), indicating that most peptide NH groups are fully exposed to solvent water in 6.0 M GdmCl. This is consistent with the previous reports that ubiquitin in concentrated denaturant (6 M GdmCl or 8 M urea) at acidic pH (pH 2.0–2.5) is almost fully unfolded (24–28). Nevertheless, several NH groups, including those of Ile3, Val5, Ile13 and Leu73, show *P* values larger than 2, and considering the error estimate of *k*_obs_, these values are significantly larger than unity. Therefore, these NH groups of ubiquitin were weakly but significantly protected against H/D exchange in 6.0 M GdmCl and pH* 2.6. Two NH groups (Ser20 and Gly75) apparently exhibited *P* values of 0.4 and 0.5, but considering the large errors of *k*_obs_ for these residues (Table 1), the *P* values might not be different from unity. The *P* values of the NH groups of the N-terminal extra residues in Table 1 were all equivalent to unity within experimental error, suggesting that this part of the protein was completely unstructured under the condition used. Therefore, we assumed that the extra-residue part did not affect the H/D-exchange behavior of the ubiquitin-sequence region of the protein.

## Discussion

We studied the H/D-exchange behavior of unfolded ubiquitin in 6.0 M GdmCl at pH* 2.6 and 20°C by the DMSO-quenched H/D-exchange method with the use of spin desalting columns for medium exchange. We successfully monitored the exchange kinetics of 27 NH groups in the ubiquitin sequence (Table 1). Although most of these NH groups were fully exposed to solvent without any significant protection against H/D exchange, the four NH groups of Ile3, Val5, Ile13 and Leu73 were significantly protected with a *P* value of 2.5–3.0, indicating the presence of residual structures in unfolded ubiquitin. In the following, we will discuss (1) native and nonnative hydrogen bonds in unfolded ubiquitin, (2) the H/D-exchange method for characterizing residual structures in unfolded states, (3) comparison with unfolded ubiquitin in 8 M urea, and (4) folding mechanisms of ubiquitin.

### Native and nonnative hydrogen bonds in unfolded ubiquitin

Because ubiquitin in 6.0 M GdmCl and pH* 2.6 was almost fully unfolded, the four weakly but significantly protected NH groups of Ile3, Val5, Ile13 and Leu73 were protected by formation of a hydrogen (H) bond with a certain acceptor group. We thus listed the H-bond lengths and the acceptor groups of the backbone NH groups in native ubiquitin (PDB code: 1UBQ) in Table 1 (12). Interestingly, three (Ile3, Val5 and Ile13) of the four protected NH groups are all H-bonded to form the first β-hairpin in the native structure; Ile3 is H-bonded with Leu15, and Val5 and Ile13 are mutually H-bonded with each other (Fig. 4) (12). This result thus strongly suggests that the first native β-hairpin, composed of residues 2–16 (Fig. 4) (12), is partially structured even in unfolded ubiquitin in 6.0 M GdmCl. The side chains of Ile3, Val5 and Ile13 are all fully buried and form a hydrophobic cluster in native ubiquitin (12), suggesting that a similar hydrophobic cluster may be organized even in 6.0 M GdmCl. In support of this statement, the formation of a native or nonnative hydrophobic cluster in concentrated denaturant (6 M GdmCl or 7–8 M urea) has also been observed in other globular proteins, including a 63-residue domain of the 434-repressor (3), lysozyme (4, 5), αlactalbumin (6) and cytochrome *ć* (7). The present results are also in agreement with the unfolded-state H/D-exchange behaviors of lysozyme (9) and acidic fibroblast growth factor (10), in which weakly protected NH groups with a *P* value of 2–3 were detected in regions highly stabilized in the native state. On the other hand, the NH group of Leu73 does not form any H-bonding interactions in the native state (12), suggesting that the Leu73 NH group may be protected by a nonnative H bonding interaction in 6.0 M GdmCl.

**FIGURE 4.**
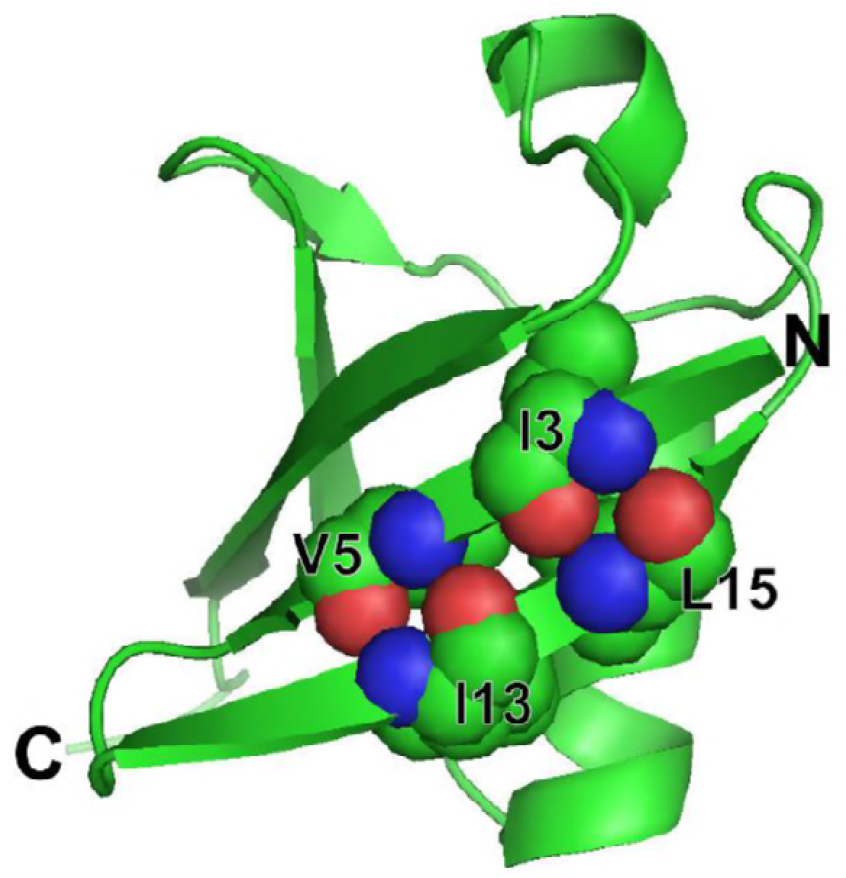
The three-dimensional structure of native ubiquitin (PDB code: 1UBQ). The four residues, Ile3, Val5, Ile13 and Leu15, in the first β-hairpin are shown in the space-filling model. The figure was drawn by PyMol (Schrödinger).

If the H/D-exchange protection is caused by formation of the H-bond, we can calculate the fraction of H-bonding (*f*_Hbond_) for the protected NH groups. Because only the non-H-bonded NH group is available for H/D exchange, (1 – *f*_Hbond_) is equivalent to *k*_obs_/*k*_int_ (= 1/*P*). Therefore, we have the following relationship: 
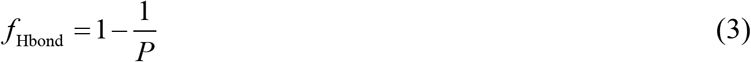

The *P* values of Ile3, Val5, Ile13 and Leu73 were 2.5–3.0, and hence the *f*_Hbond_ values of the NH groups of these residues were estimated at 0.60–0.67, indicating that the NH groups were 60–67% H-bonded in unfolded ubiquitin in 6.0 M GdmCl at pH* 2.6 and 20°C.

### The H/D-exchange method for characterizing residual structures in unfolded states

We used the DMSO-quenched H/D-exchange method to characterize the residual structure in unfolded ubiquitin in 6.0 M GdmCl at pH* 2.6 and 20°C. We also utilized spin desalting columns for the medium exchange from 6.0 M GdmCl to the quenching DMSO solution (95% DMSO-*d*_6_/5% D_2_O, pH* 5.0) (11). Because the H/D-exchange reaction in the unfolded protein takes place rapidly with a *k*obs value close to *k*_int_, we have to quench the exchange reaction, after a predetermined exchange time, for the subsequent analysis of the NH-proton signals by NMR. The use of spin desalting columns was very effective for this purpose, and we could realize a quick medium exchange from 6.0 M GdmCl to the quenching solution (11). As a consequence, we could successfully observe the weakly but significantly protected NH protons, which were protected not only by the native H-bonding interactions (Ile3, Val5 and Ile13) but also by the non-native H-bonding interaction (Leu73). In previous H/D-exchange studies on unfolded proteins, rapid refolding to the native state was used for quenching the H/D-exchange reaction in the unfolded state (9, 10). However, when rapid refolding is used for the quenching, only the NH protons protected in the native state are available for detection. Therefore, the NH proton of Leu73 of ubiquitin, which was not protected in the native state (29), could not be detected by the use of rapid refolding, indicating that spin desalting columns would be effective for studies of the unfolded-state H/D-exchange.

### Comparison with unfolded ubiquitin in 8 M urea

Ubiquitin in 8 M urea at acidic pH (pH 2.0–2.5) has been used as a model of unfolded proteins for more than a decade now, and studied by a variety of physicochemical techniques, including hetero-nuclear NMR spectroscopy (26–28) (30, 31), small angle X-ray scattering (SAXS) (32), and molecular dynamics (MD) simulations (33). However, the results thus far reported have been rather controversial.

Schwalbe and his coworkers (26–28) measured ^3^*J* coupling constants, dipole–dipole cross correlation relaxation rates, and ^15^N chemical shifts and relaxation rates by hetero-nuclear NMR spectroscopy, and concluded that ubiquitin in 8 M urea at pH 2 was completely unfolded and devoid of residual non-random structure.

On the other hand, Grzesiek and his coworkers (30) studied highly sensitive H^N^–N^N^ residual dipolar couplings of unfolded perdeuterated ubiquitin in 8 M urea at pH 2.5, and reported evidence of the persistence of native-like structure in ubiquitin’s first β-hairpin. They also measured chemical shifts, ^3^*J*HNHA couplings, relaxation rates, and ^h3^*J*NC’ couplings to directly detect H-bonds, and concluded that the native first β-hairpin conformation was populated to at most 25% in unfolded ubiquitin in 8 M urea. Structural ensembles of unfolded ubiquitin in 8 M urea, calculated by the program X-PLOR-NIH as well as by MD simulations with the restraints of the NMR and SAXS data (31–33), also revealed significant (10–20%) population of the native-like first β-hairpin and nonnative α-helical conformation in the C-terminal half, and hence, urea-unfolded ubiquitin has some similarities to its methanol/acid-unfolded A state (34). These results are thus fully consistent with the results in the present study. We observed that the first native β-hairpin was partially structured and that the NH group of Leu73 underwent a nonnative H-bonding interaction; the latter result was likely due to the formation of the nonnative α-helix in the C-terminal half of unfolded ubiquitin. The protection pattern of NH groups observed here thus shows a certain similarity to the pattern previously reported for the A state (29), although the *P* values in the A state are approximately ten-fold higher than those in unfolded ubiquitin in 6.0 M GdmCl.

The present study used the DMSO-quenched H/D exchange method, which is completely different from the above mentioned various sophisticated NMR methods, SAXS and MD simulations, but nevertheless, we observed the same residual structures in unfolded ubiquitin in 6.0 M GdmCl, providing strong evidence that unfolded ubiquitin in a concentrated denaturant (6 M GdmCl or 8 M urea) at acidic pH possesses residual structures: the partially structured first native β-hairpin and the nonnative H-bonding interaction near the C-terminus. The present results also showed that the partially structured β-hairpin was not uniformly organized, i.e., only the three NH groups of Ile3, Val5 and Ile13 had a *f*Hbond value of 0.6–0.67, but almost all the other NH groups involved in the first β-hairpin (residues 2–16) were not protected against H/D-exchange (Table 1).

### Folding mechanisms of ubiquitin

The presence of the residual structures, i.e., the native-like first β-hairpin and the nonnative H-bonding interaction near the C-terminus in unfolded ubiquitin in 6.0 M GdmCl, strongly suggests that they may play significant roles in the refolding of the protein from the unfolded state. Real refolding experiments of globular proteins usually begin with the unfolded state in concentrated denaturant (6 M GdmCl or 8 M urea), and the residual structures present, if any, in the unfolded state may control the folding mechanisms of the proteins.

Using pulsed H/D-exchange with rapid mixing methods and 2D NMR analysis, Briggs and Roder (13) studied early H-bonding events in the ubiquitin refolding from the GdmCl-unfolded state, and reported that the NH protons in the first β-hairpin and an α-helix of residues 23–34 become 80% protected in an initial 8-ms folding phase. Sosnick and his co-workers (14, 15) used mutational ϕ-value analysis as well as ψ-analysis, which utilized engineered bi-His metal binding sites, to investigate the transition state of the ubiquitin refolding from the GdmCl-unfolded state. The transition state thus characterized had the partially folded first β-hairpin and α-helix. The transition-state structure of ubiquitin was also characterized by Went and Jackson (16) using a comprehensive ϕ-value analysis, and the structure was highly polarized, with medium and high ϕ-values found only in the N-terminal region of the protein, including the first β-hairpin. Therefore, in all these previous studies on ubiquitin folding, the first β-hairpin was significantly structured in the folding intermediate and the transition state, while the C-terminal region of the protein was less organized. It is thus reasonable to conclude that the residual native β-hairpin in unfolded ubiquitin in 6 M GdmCl forms a folding initiation site and guides the subsequent folding reactions of the protein.

## Conclusions

The residual structures in unfolded ubiquitin in 6.0 M GdmCl at pH* 2.6 were detected and characterized by the DMSO-quenched H/D-exchange method with the use of spin desalting columns for the medium exchange. By the resonance assignment of the NH proton signals in the ^1^H– ^15^N HSQC spectrum of the protein dissolved in the quenching DMSO solution, we successfully monitored the H/D-exchange kinetics of the 27 NH groups in the ubiquitin sequence. Although most of these NH groups were almost fully exposed to solvent with a *P* value close to unity, the four NH groups of Ile3, Val5, Ile13 and Leu73 were weakly but significantly protected with a *P* value of 2.5–3.0. The results strongly suggest that the first native β-hairpin, composed of residues 2–16, is partially structured even in 6.0 M GdmCl and that the NH group of Leu73 is protected by a nonnative H-bonding interaction in the unfolded state. From comparison of the present results with the previous studies on the folding kinetics of ubiquitin, it is concluded that the residual native β-hairpin in unfolded ubiquitin in 6 M GdmCl forms a folding initiation site and guides the subsequent folding reactions of the protein.

## Author Contributions

M.S.C., T.N., K.Kato and K.Kuwajima designed the research. M.S.C., T.N. and M.Y.-U. performed experiments. M.S.C., T.N., M.Y.-U., T.Y. and K.Kuwajima analyzed the data. M.S.C. and K.Kuwajima wrote the article.

## Acknowledgments

We thank Ms. Ying-Hui Wang (Okazaki Institute for Integrative Bioscience) for her help in spectral analyses. This study was supported in part by MEXT KAKENHI Grant Numbers JP20107009, JP25102001 and JP25102008, by JSPS KAKENHI Grant Numbers JP25440075, JP15H02491, JP16K07314 and JP17K15441, by the Nanotechnology Platform Project of the Institute for Molecular Science, and by the Okazaki ORION project of the Okazaki Institute for Integrative Bioscience.

